# Meta-analysis of heat-stressed transcriptomes using the public gene expression database from human and mouse samples

**DOI:** 10.1101/2023.04.30.537112

**Authors:** Sora Yonezawa, Hidemasa Bono

**Author notes:** Corresponding author: Hidemasa Bono.

## Abstract

**Background:** Climate change has significantly increased the frequency of exposure to heat, adversely affecting human health and various industrial sectors. Heat stress is an environmental stress defined as the exposure of organisms and cells to abnormally high temperatures. Heat-stress research has predominantly focused on response systems involving heat shock factors acting as transcription factors and heat shock proteins functioning as molecular chaperones. However, to comprehensively elucidate the mechanisms underlying an organism’s response to heat stress, it is essential to investigate and analyze genes that have been underrepresented, less well-known, or overlooked in previous studies. In this study, we analyzed heat stress-responsive genes using a meta-analysis of numerous gene expression datasets.

**Results:** First, we collected paired heat exposure and control data from public databases. Gene expression data were obtained for 322 human and 242 mouse pairs. The expression ratios (HN-ratios) of the collected pairs were calculated, and the identification of upregulated and downregulated expression profiles was determined according to defined thresholds. The number of upregulated and downregulated genes was calculated as the heat stress - non-treatment score (HN-score), which is the value of: [number of upregulated genes] - [number of downregulated genes] for each gene and was used as the index of analysis. The HN-score comprehensively evaluated gene expression variation, and 76 upregulated and 37 downregulated genes common to human and mouse were identified. We performed enrichment, protein-protein interaction network, and transcription factor target gene analyses. Furthermore, we evaluated the extracted genes through integrated analysis using publicly available ChIP-seq data for HSF1, HSF2, and PPARGC1A (PGC1-α), and gene2pubmed data, which were sourced from previous literature. The results identified previously overlooked genes, such as *ABHD3*, *ZFAND2A*, and *USPL1*, as commonly upregulated genes.

**Conclusions:** Based on the findings of this study, further functional analysis of the extracted genes using genome editing and other technologies has the potential to contribute to coping with climate change and potentially lead to new knowledge and technological advances.

## Background

Climate change is expected to result in higher average temperatures and an increased intensity and frequency of heat waves, thereby increasing the adverse effects of high temperature. For example, it has been reported that the average temperature on land where humans reside was projected to be 0–6 °C higher in summer of 2021 than that in the years 1986–2005. Moreover, an increase in the number of heat-wave days experienced by children under the age of 1-year and adults over the age of 65 years, which are age groups vulnerable to high temperatures, has been observed. These phenomena are anticipated to cause heat-related diseases (e.g., heat stroke) and are considered as indicators of the adverse effects of climate change on human health[1–3]. Visual climate change data are available at https://www.lancetcountdown.org/data-platform/.

High-temperature environments impose stress on organisms and cause various cellular changes. Heat-stress (HS, also known as heat shock or hyperthermia) is an environmental stress that occurs when organisms and cells are exposed to abnormally high temperatures, which are thought to increase with climate change. HS has diverse effects on the elements that form cells[4]. At the cellular level, it causes the uncoupling of oxidative phosphorylation in mitochondria, a cellular organelle, and promotes the generation of reactive oxygen species (ROS)[5,6]. Additionally, it causes physical changes in the biomembrane, such as changes in membrane fluidity[7]. HS exerts various effects at the molecular level; for example, it can inhibit DNA repair pathways and cause DNA damage[8–10]. HS can also denature intracellular proteins and lead to their aggregation[11].

A programmed gene expression response called the heat shock response (HSR) is a cellular response to HS. The HSR is primarily regulated by a group of transcription factors called heat shock factors (HSFs); of them, HSF1 plays a central role in the response. When triggered by HS, HSF1 forms a trimer and binds to heat shock elements (HSEs) in DNA, which induces the expression of genes encoding heat shock proteins (HSPs). The main HSP members are important proteins that function as molecular chaperones and maintain protein homeostasis (proteostasis) by interacting with other proteins, stabilizing them, and helping them to acquire functional conformations. These members are mainly classified as HSP40, HSP60, HSP70, HSP90, HSP100, and sHSPs (small HSPs), and they allow cells to maintain proteostasis and survive[12–14].

To date, HS-related studies have primarily focused on proteins that function as molecular chaperones, accumulating knowledge regarding their role in maintaining cell proteostasis and protecting cells under stress conditions, including heat stress. However, heat affects numerous cellular components [4]; moreover, proteins induced by heat stress have been proposed to be classified into functional classes such as nucleic acids and metabolism [12]. Despite this classification, there have been limited available studies on the genes encoding these proteins induced by HS. Therefore, a comprehensive analysis, including previously uncharacterized or poorly known HS-related genes, is essential to understand the heat stress response deeply. One approach to solving this problem is data-driven research using meta-analysis. Meta-analyses of gene expression data related to hypoxia and oxidative stress have been conducted in human, mouse, and both species, yielding novel findings, including discovering genes overlooked in previous hypothesis-driven studies [15–17].

This study aimed to analyze genes involved in HS by collecting HS-related gene expression data from public databases in human and mouse and perform a meta-analysis. As a data-driven study, meta-analysis provides an approach different from that used in previous hypothesis-driven studies and may yield novel insights. The dataset collected in this study and the selected gene set will contribute to efforts to reveal a more comprehensive understanding of organisms’ responses to HS.

## Results

### Characteristics of HS-related high-throughput sequencing data

In this study, we obtained 66 Sequence Read Archive (SRA) IDs and a total of 564 pairs of gene expression data. The gene expression data consisted of 322 pairs for human and 242 pairs for mouse, which compared HS and non-treatment conditions. We searched for RNA sequencing (RNA-seq) data, but encountered reports using different sequencing methods. Therefore, to obtain a broader dataset, we collected gene expression data using different methods, such as RNA-seq and Ribo-seq (also known as ribosome profiling). A summary of the sample types, temperature conditions, and treatment times for the collected data is shown in Fig. 1. In human samples, primarily cultured cells were collected, while in mouse samples, both cultured cells and tissues were collected. The samples with the most data included cultured human cells (MRC5-VA: 55 pairs), mouse cells (MEF cells: 52 pairs), and mouse tissues (kidneys: 23 pairs) (Fig. 1A). In human samples, 172 pairs (30.5%) were derived from cancer cells. The most common conditions were a temperature of 42 °C for human and mouse samples, and a treatment time of 60 min for human samples and 30 min for mouse samples (Fig. 1B, C). The conditions included in the “other” category comprised samples with discontinuous treatment times, such as a sample in which the temperature was changed midway from 42 °C to 48 °C, or samples exposed to high-temperature conditions repeatedly for 3 h per day for over 1 week (Fig. 1B, C). Metadata, including detailed information on temperature conditions, sample types, the sequencing type, is available from the figshare file (Table S1, 2) [18].

**Fig. 1.**
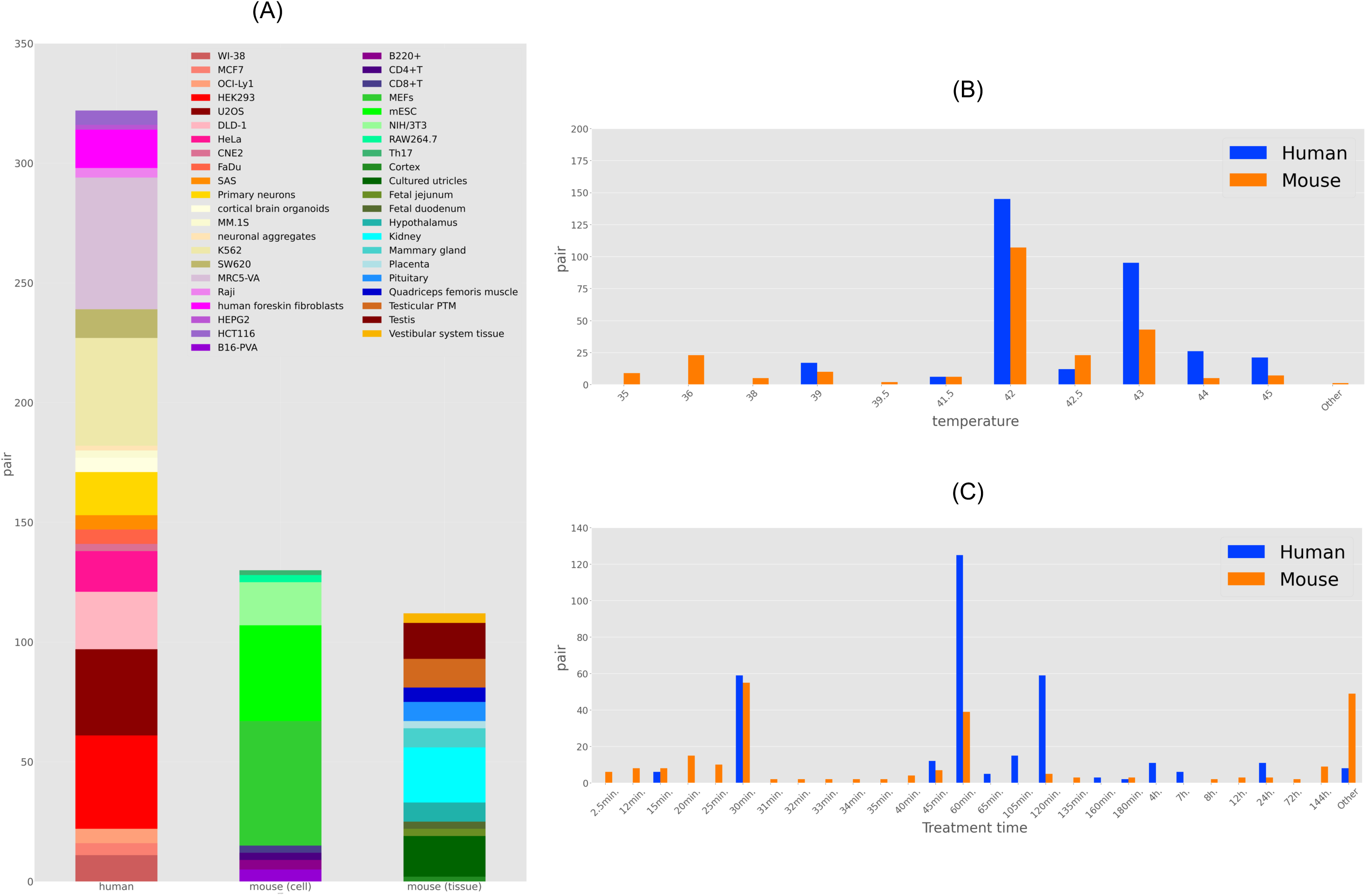
Summary of the contents of heat stress-related dataset. (A) The stacked bar graph shows the number of data pairs collected from the sample. From left to right: human cell, mouse cell, and mouse tissue types. (B, C) The dataset summary for (B) temperature condition (degree, °C), and (C) treatment time.

### Verification of upregulated and downregulated genes in each organism

The HN-ratios (expression ratios) and HN-scores (as an index of analysis) were calculated for all genes listed in GENCODE Release 37 (GRCh38.p13) for human, and GENCODE Release 26 (GRCm39) for mouse[16,17]; the calculations were performed for meta-analysis purposes of gene expression data from heat-exposed human and mouse samples. The HN-score for each gene represented the difference between the number of experiments with increased expression results and those with decreased expression results. Scatter plots of the HN-scores calculated at 5-fold or 1/5-fold expression ratio thresholds for all genes in human and mouse, respectively, are shown in Fig. 2A and D. The top 500 genes with high HN-scores and the bottom 500 genes with low HN-scores were extracted from human and mouse samples, respectively. The top 500 genes were designated as the “upregulated gene group,” while the bottom 500 genes were referred to as the “downregulated gene group.” These gene sets were used for the subsequent analyses.

**Fig. 2.**
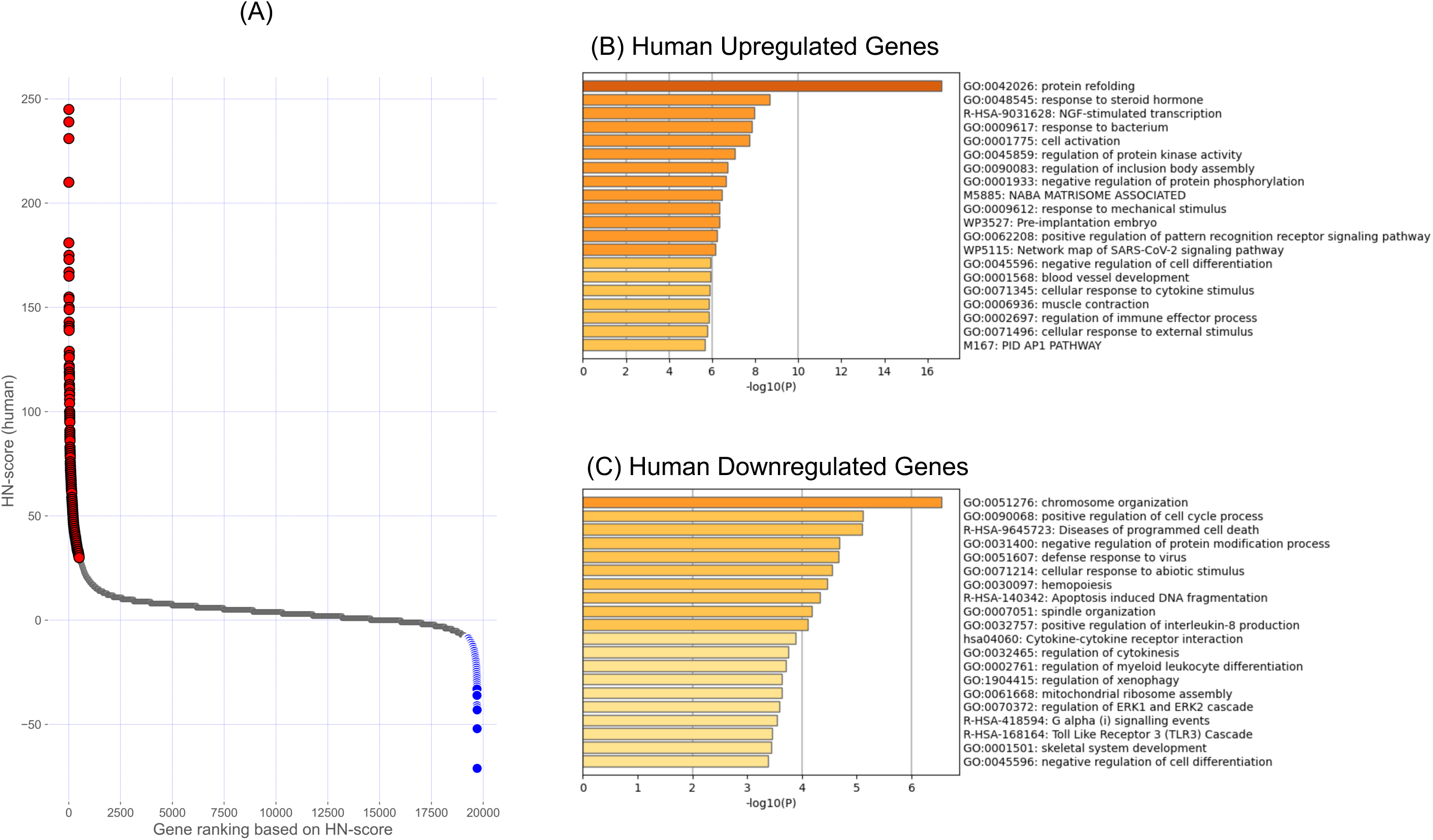
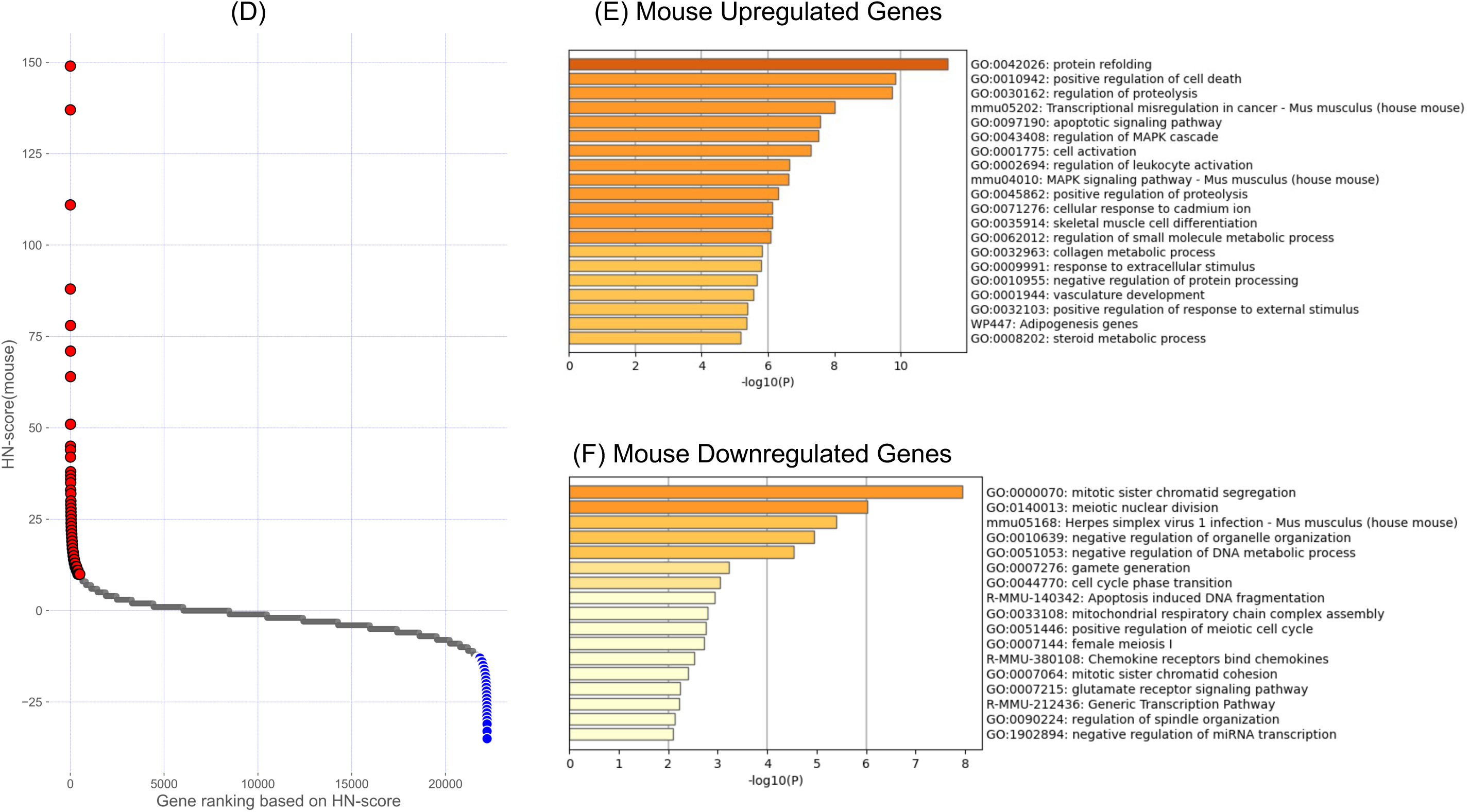
Scatter plots of the HN-score and gene set enrichment analysis of upregulated and downregulated genes in human and mouse. (A, D) Scatter plots of HN-scores for all genes in (A) human and (D) mouse. Red and blue dots represent the 500 genes with high and low HN-scores, respectively. (B, E) Results of gene set enrichment analysis of upregulated genes in (B) human and (E) mouse. (C, F) Gene set enrichment analysis of the downregulated genes in (C) human and (F) mouse.

Gene set enrichment analysis was performed to characterize the upregulated and downregulated gene groups in human and mouse. In the upregulated gene group, “protein folding” (Gene Ontology (GO):0042026) was the most significantly enriched GO term in human and mouse (Fig. 2B, E). In contrast, in the downregulated gene group, “chromosome organization” (GO:0051276) was the most enriched GO term in human samples, while “mitotic sister chromatid segregation” (GO:0000070) was the most enriched term in mouse samples (Fig. 2C, F).

### Identification of common upregulated or downregulated genes in human and mouse

From the obtained upregulated and downregulated gene sets in human and mouse, we extracted sets of genes common to both human and mouse samples to analyze the genes with a typical response to HS. We accessed Mouse Genome Informatics (MGI)[19], a database that compiled a variety of information on mouse, and used the Batch Query function to convert the gene symbols of the human upregulated and downregulated gene sets into the corresponding mouse ortholog gene symbols denoted by “Current Gene Symbol” (e.g., *HSPA1A* → *Hspa1a*). Consequently, 410 upregulated genes and 406 downregulated genes from the human set were converted into mouse orthologs. Of them, 76 genes were commonly upregulated, and 37 were commonly downregulated (Fig. 3A, B). A scatter plot was created to evaluate the distribution of the HN-scores for the common genes using the HN-scores of the corresponding orthologous genes in human and mouse (Fig. 3C). In the scatter plot, the 76 commonly upregulated genes are shown in red, and the 37 commonly downregulated genes are shown in blue. As shown in Fig. 3C, a notable pattern was observed for *HSPA1A* and *HSPA1B*, which encode the HSP70 family of proteins. Various stressors, including HS, induce expression of these genes. Using the ShinyGO(ver. 0.77)[20], the common upregulated and downregulated genes were plotted against the human genome, revealing their distribution (figshare Fig. S1). [18]. Among the 76 common upregulated genes, 10 genes, including *HSPA1A* and *HSPA1B*, were annotated as “response to heat” (GO:0009408). The list of upregulated and downregulated genes with symbols and HN-score data for human and mouse is available in figshare (Tables S3 and S4)[18]. Additionally, a sample breakdown of the HN-scores for common upregulated and downregulated genes is shown on figshare (Fig. S2)[18].

**Fig 3.**
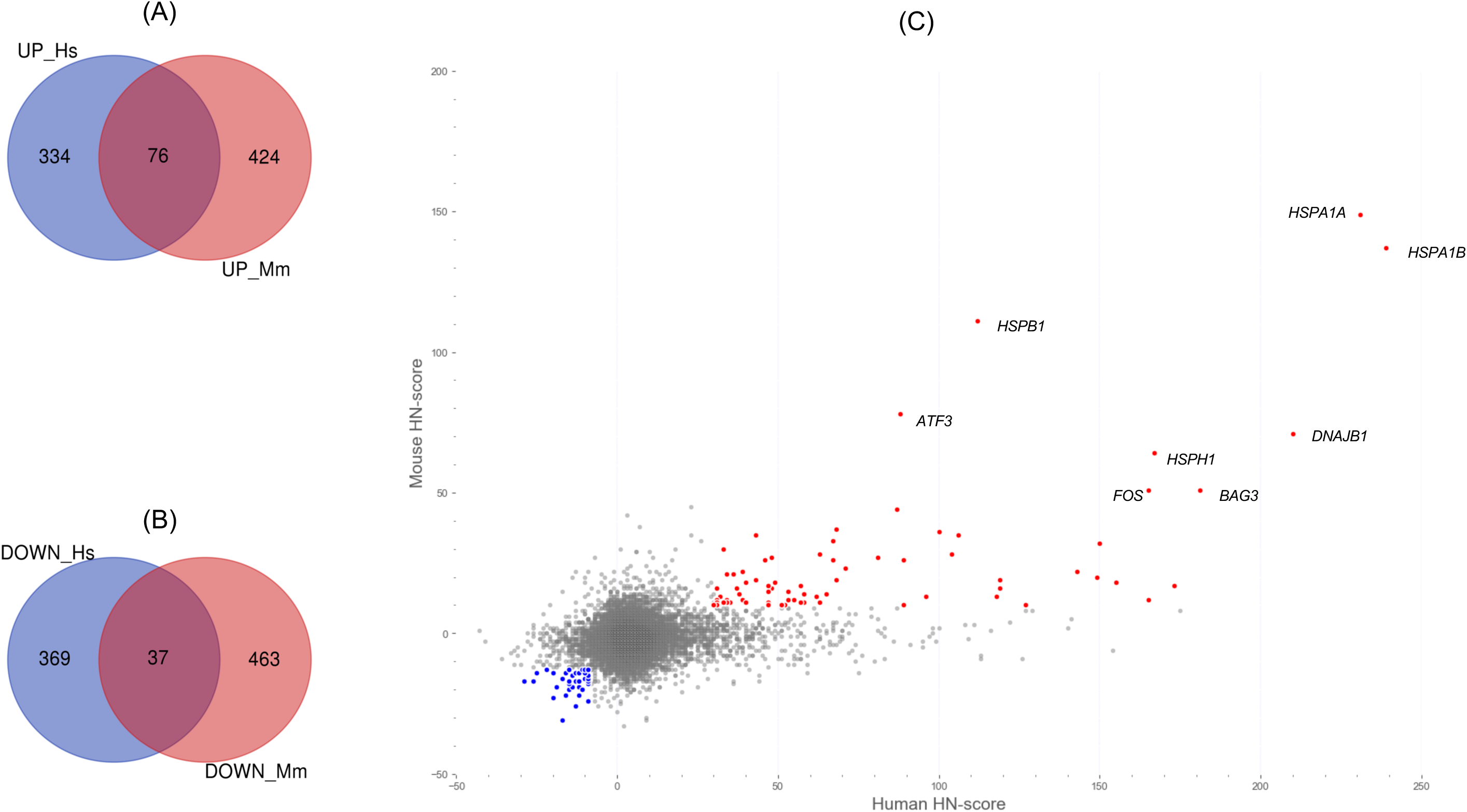
Overlaps of 76 upregulated genes and 37 downregulated genes in human and mouse using mouse gene symbols. (A) Venn diagram of the upregulated genes in human (UP_Hs) and mouse (UP_Mm). (B) Venn diagram of downregulated genes in human (DOWN_Hs) and mouse (DOWN_Mm). (C) Scatter plots of common up/downregulated genes in human and mouse. The x-axis represents the human HN-score, and the y-axis represents the mouse HN-score. The red plot indicates the 76 commonly upregulated genes, whereas the blue plot indicates the 37 commonly downregulated genes, with the upper right indicating a higher HN-score and the lower left indicating a lower HN-score. The gene symbols shown in the figure indicate genes with HN-scores of 50 or higher in human and mouse, respectively.

We performed gene set enrichment analysis with human gene symbols using Metascape, as previously conducted in the “*Verification of upregulated and downregulated genes in each organism*” section, to characterize the 76 common upregulated genes and 37 common downregulated genes (Fig. 4A, B). Additionally, the search results using mouse gene symbols are shown in figshare (Fig. S3). Among the common upregulated genes, “response to topologically incorrect protein” (GO:0035966) was the most enriched (Fig. 4A). Within this term, the following genes were found: (1) Heat shock 70kDa proteins (HSPA): *HSPA1A*, *HSPA1B*, *HSPA1L*, *HSPA4L*, *HSPA8*, *HSPH1*; (2) DNAJ (HSP40) heat shock proteins: *DNAJA1*, *DNAJA4*, *DNAJB1*, *DNAJB4*; (3) small heat shock proteins (HSPB): *HSPB1*, *HSPB8*; (4) Chaperonins: *HSPD1*, *HSPE1*; (5) Heat shock 90kDa proteins (HSP90): *HSP90AA1*; (6) BAG cochaperones (BAG): *BAG3*; and (7) Serpin peptide inhibitors (SERPIN): *SERPINH1* (also known as hsp47). Thus, the term covers genes encoding various classes of molecular chaperones and cofactors, and these genes, which play a role in maintaining proteostasis, are commonly expressed in both human and mouse. Alternatively, the term “negative regulation of chromosome organization” (GO:2001251) was found to be the most enriched among the common downregulated genes (Fig. 4B). This term includes genes that play essential roles in the regulation of mitosis, such as *PLK1* and *HASPIN*.

**Fig 4.**
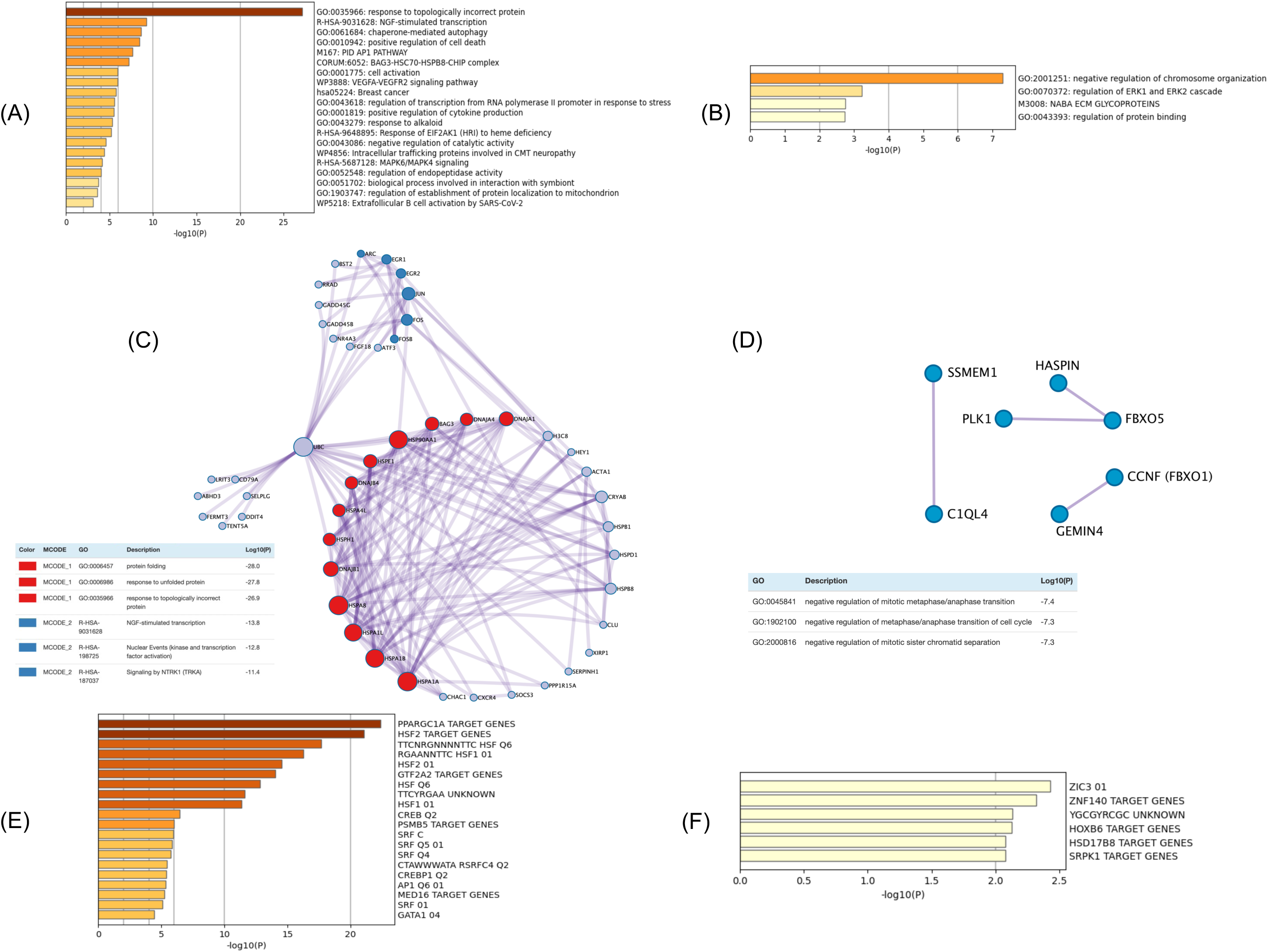
Various functional analysis of common upregulated/downregulated genes. (A, B) gene set enrichment analysis of common (A) upregulated genes and (B) downregulated genes. (C, D) protein-protein interaction network analysis of common (C) upregulated genes and (D) downregulated genes. (E, F) transcription factor target enrichment analysis of common (E) upregulated genes and (F) downregulated genes.

Protein–protein interaction networks were analyzed for more detailed information on the commonly upregulated and downregulated genes annotated in the enriched GO terms (Fig. 4C, D). For common upregulated genes, a cluster of genes encoding molecular chaperones was visualized using the Molecular Complex Detection (MCODE) algorithm (Fig. 4C; shown in red). Groups of genes classified as immediate early genes (IEGs), including *FOS*, *FOSB,* and *JUN*, which encode transcription factor activator protein 1 (AP-1), were also clustered (Fig. 4C; shown in blue). IEGs are a group of genes rapidly expressed in response to cellular stimuli. The *FOS* and *JUN* genes extracted in this study have been well-characterized in previous studies. *EGR1*, *EGR2*, *FOS*, *FOSB*, and *ARC* of IEGs in this blue cluster were annotated in the “NGF-stimulated transcription” pathway (R-HSA-9031628). Additionally, actin alpha 1 skeletal muscle (*ACTA1*), which belongs to the actin family, was also found in the protein–protein interaction network, although it was not included in the cluster. In contrast, no closely linked networks were formed for the downregulated genes compared to those observed for the upregulated genes (Fig. 4D).

We conducted enrichment analysis to obtain an overview of the gene expression regulatory information for common upregulated genes and common downregulated genes (Fig. 4E, F). Gene sets were obtained from the Molecular Signatures Database (MSigDB)[21]. Among the commonly upregulated genes, PPARGC1A_TARGET_GENES (systematic name M30124) was the most enriched (Fig. 4E). *PPARGC1A* encodes peroxisome proliferator-activated receptor gamma coactivator 1α (PGC1-α), a critical regulator of mitochondrial biogenesis. Terms related to HSF1 and HSF2, the central regulators of HS, such as HSF2_TARGET_GENES (systematic name: M30020), TTCNRGNNNNTTC_HSF_Q6 (systematic name: M16482), and RGAANNTTC_HSF1_01 (systematic name: M8746), were also included. Genes common to these four sets, such as *HSPA1A*, which encodes a molecular chaperone, were also included. The common genes are available on figshare (Fig. S4). Based on these results, we conducted a detailed analysis on HSF1, HSF2, and PPARGC1A. A gene set related to serum response factor (SRF), SRF_C (systematic name: M12443), was also found in the enrichment analysis, although it was not as enriched (Fig. 4E). This gene set contained IEGs (*FOS*, *FOSB*, *EGR1*, and *EGR2*) included in the blue cluster shown in Fig. 4C, and *ACTA1*, which was not included in the blue cluster. However, in the common downregulated genes, enrichment analysis results were inconclusive (Fig. 4F).

### Integration of HN-score, transcription factor-binding information, and literature information

Transcription factor target gene enrichment analysis revealed that the target genes of HSF1, HSF2, and PPARGC1A were enriched in the common upregulated genes (Fig. 4E). To further analyze the binding information of these three transcription factors, we integrated ChIP-seq data from the ChIP-Atlas database[22], processed using the MACS2 program, with the HN-score results obtained in this study. Scatter plots illustrating ChIP-seq and HN-scores were used for visualization (Fig. 5). Additionally, we used gene2pubmed data to visualize the number of reports registered in PubMed for each of the 76 common upregulated genes. The source data can be obtained from figshare (Table S5). As shown in Fig. 5A, in addition to HSP genes such as *HSPA1A*, which are known to be targets of HSF1, abhydrolase domain-containing 3 (*ABHD3*) was identified as a target candidate among genes with fewer research reports. In contrast, genes with low ChIP-seq scores were observed. *EGR1* and *ACTA1*, the target genes of SRF (Fig. 4C, E), showed such characteristics, suggesting that they belong to pathways unrelated to HSF1. Common genes with high ChIP-seq scores were found not only for HSF1 (Fig. 5A), but also for HSF2 (Fig. 5B) and PPARGC1A (Fig. 5C). In addition to the HSP genes, zinc finger AN1-type containing 2A (*ZFAND2A*) and ubiquitin-specific peptidase-like 1 (*USPL1*), had high ChIP-seq scores (Fig. 5).

**Fig 5.**
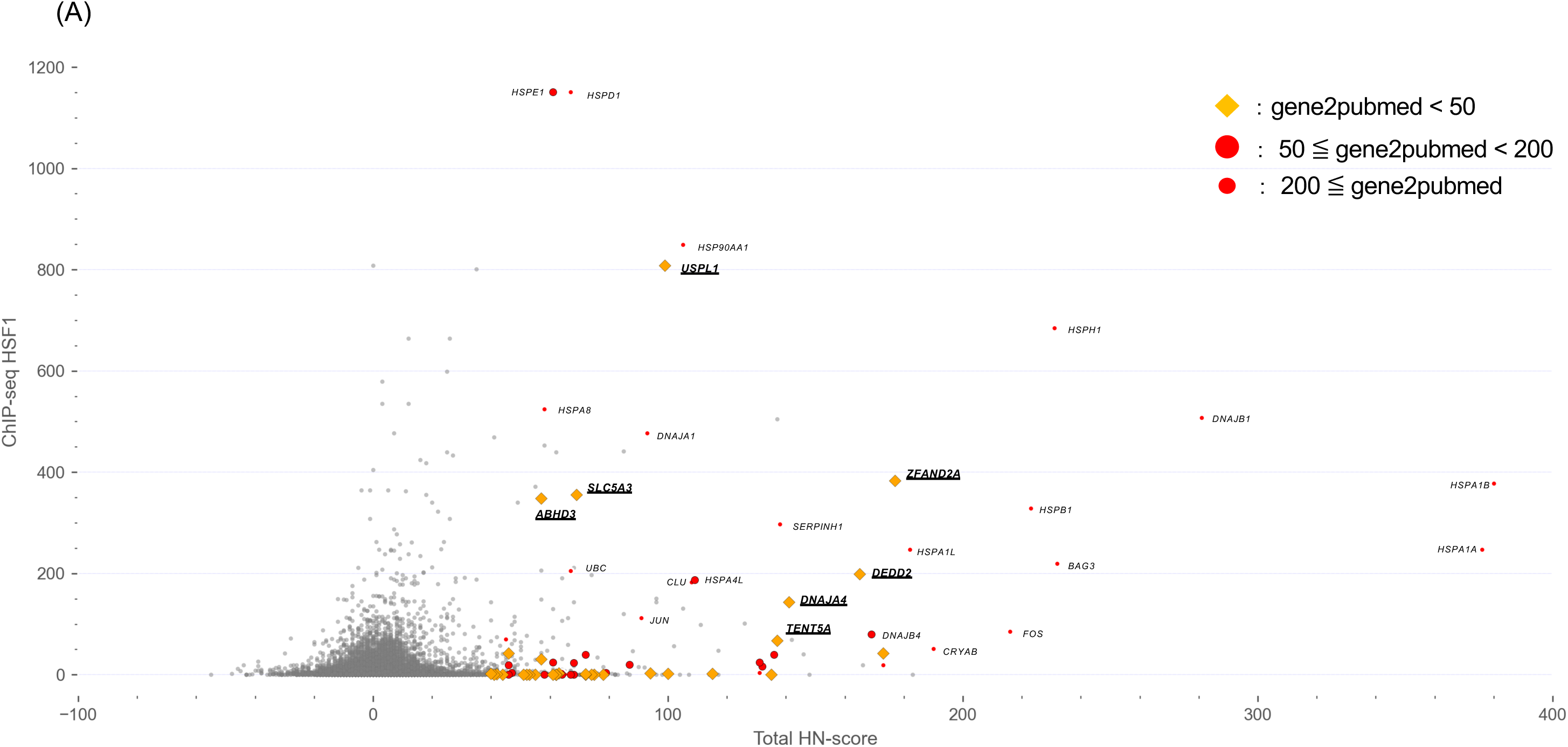
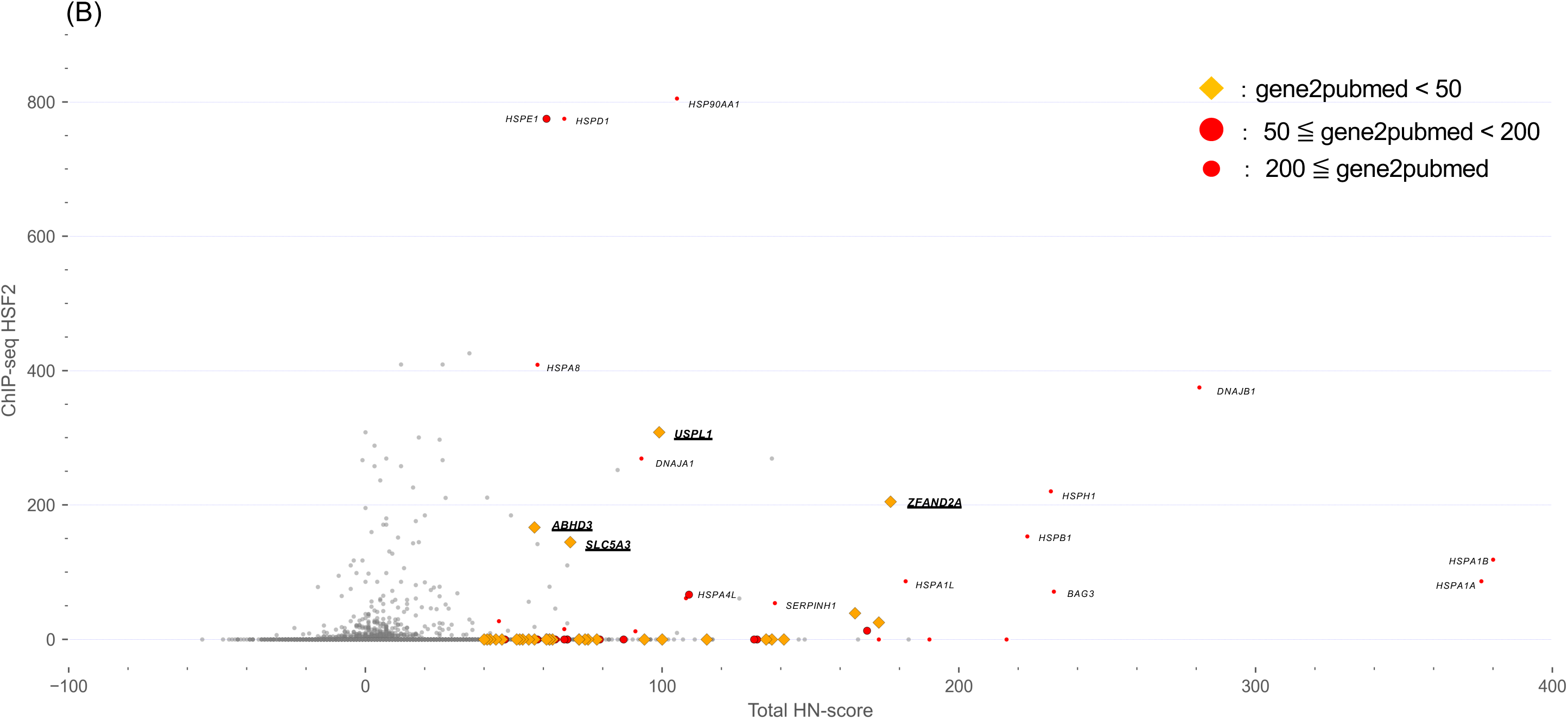
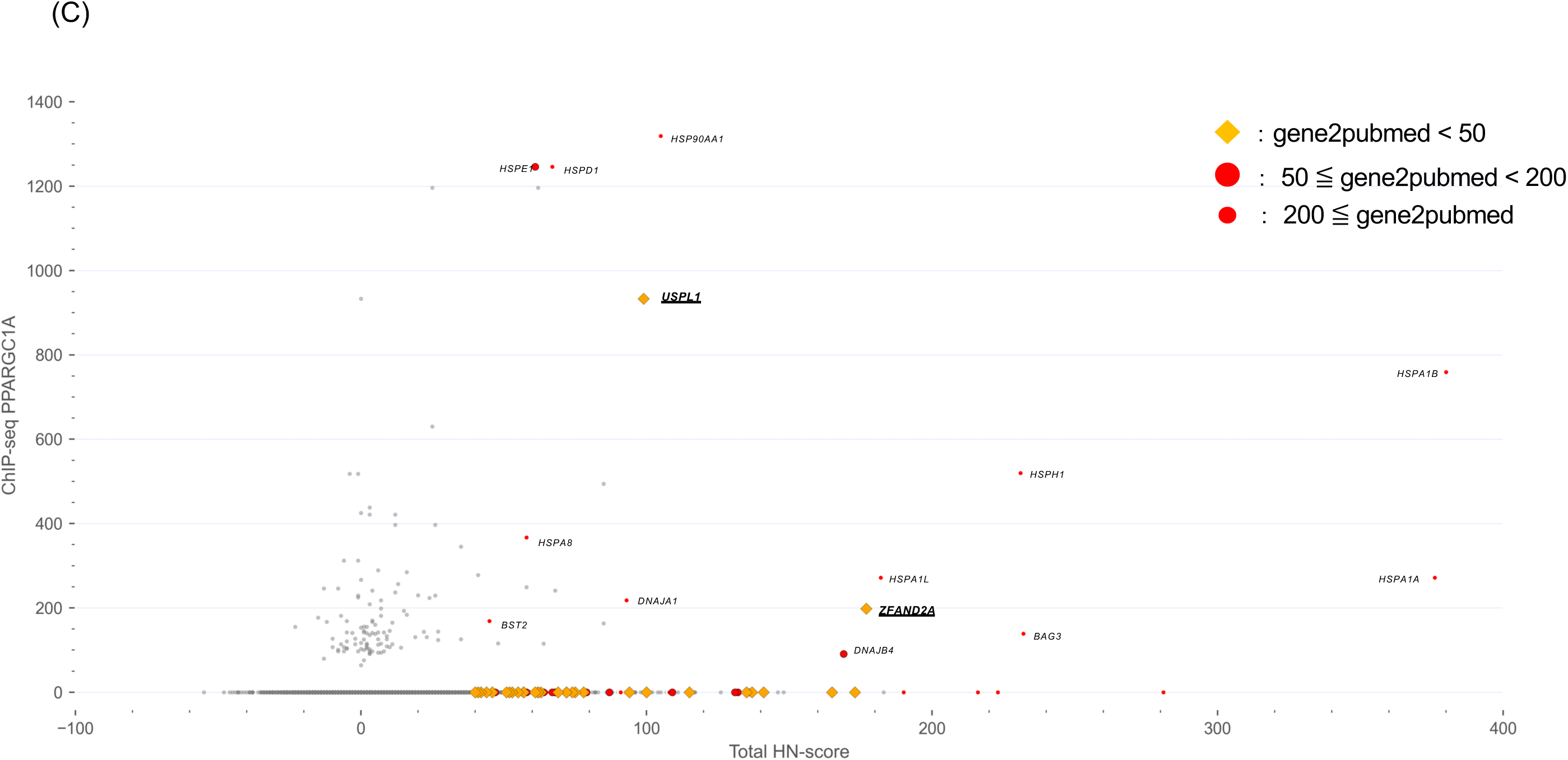
Integrated scatter plots using HN-score, average MACS2 score, and gene2pubmed data for several transcription factors. (A, B, C) Scatter plots from the integration of the HN-score, ChIP-seq data, and gene2pubmed data. X-axis “Total HN-score” means (human HN-score) + (mouse HN-score). Y-axis indicates the average MACS2 score (peak mean) for the transcription factors (A) HSF1, (B) HSF2, and (C) PPARGC1A. Average MACS2 scores were calculated from human information. ChIP-seq data counts are (A) HSF1:105, (B) HSF2:25, and (C) PPARGC1A:1. Using gene2pubmed data, yellow diamonds were plotted against common upregulated genes if the number of reported papers was less than 50, large red circles if the number of reported papers was between 50 and 200, and small red circles if the number of reported papers was 200 or more.

## Discussion

In this study, we manually curated 322 human and 242 mouse pairs of heat-exposed and untreated sample data from public databases for gene expression data, and performed HS response gene analysis. The top 500 genes with the highest HN-scores in human and mouse were selected as upregulated genes, and the bottom 500 were selected as downregulated genes for gene set enrichment analysis. GO terms associated with protein folding were the most enriched in the upregulated gene dataset for both species (Fig. 2A, D). Considering that HS disrupts proteostasis[4,12], this gene set reflects the variation in gene expression under HS conditions reported in previous studies, indicating that HS- related genes were properly extracted.

To identify common HS response mechanisms, we narrowed down our results to the upregulated and downregulated genes common to both human and mouse and identified 76 upregulated genes and 37 downregulated genes (Fig. 3A, B). Detailed information on these two gene sets was obtained by protein–protein interaction network analysis, transcription factor target gene enrichment analysis, and gene set enrichment analysis (Fig. 4). Several genes encoding molecular chaperones, including *HSPA1A* and *HSPA1B*, were included in the common upregulated genes and clusters of these genes were created (Fig. 4A, C). Results from meta-analysis have also revealed this response to proteostasis induced by HS. In contrast, genes associated with different processes, such as IEGs and *ACTA1*, were also present in the common upregulated genes (Fig. 4C). These genes are also targets of the transcription factor SRF, which is distinct from HSF1, the master regulator of the HSR (Fig. 4E). It is already known that SRF is associated with the expression of IEGs (e.g., *FOS*)[23]. In addition, genome-wide analysis has revealed that SRF mainly regulates genes induced early in HS in an HSF1-independent manner, and many of these genes are related to the cytoskeleton[24]. Therefore, genes associated with HSF1-independent pathways were identified.

Moreover, in addition to the HN-score data for each gene calculated in this study, ChIP-seq data as transcription factor-binding information and gene2pubmed data as literature information were integrated to visualize the characteristics of each gene, and information on common upregulated genes in human and mouse was added (Fig. 5).

In the ChIP-Atlas database, registered ChIP-seq data were scored using the MACS2 score program, with higher scores suggesting direct binding of transcription factors[22]. This information can be used to filter genes that are directly or indirectly regulated by the transcription factors of interest[15]. As shown in Fig. 5A, genes directly regulated by HSF1, the master regulator of the response to HS, and genes thought to be indirectly regulated were included in the common upregulated genes. In addition to HSF1, we targeted HSF2, a member of the HSF family, and PPARGC1A (PGC1-α), which is associated with various metabolic events because of the enrichment of genes encoding different transcription factors (Fig. 4E). HSF2 interacts with HSF1, forming a heterodimer with HSF1 under HS conditions [13,25]. On the contrary, PPARGC1A (PGC1-α) may be associated with the induction of the expression of representative HSP genes in the red cluster in Fig. 4C, and has been shown to activate *Hspa1a* transcription by interacting with HSF1 in mouse 10T1/2 cells [26]. Thus, the regulation of gene expression in response to HS may be coordinated with transcription factors other than HSF1. The results of this study also suggest that this is a common gene with a high ChIP-seq score, as shown in Fig. 5. Thus, the integration and use of transcription factor-binding information suggests that it may be a helpful method for a more detailed analysis of the response to HS.

In addition to information on transcription factor regulation using ChIP-seq data, gene2pubmed data were used to assess gene attention and visualize the results (Fig. 5). We investigated 76 common upregulated genes with high ChIP-seq scores. As shown in Fig. 5A, *ABHD3*, with a high ChIP-seq score, was implicated in the degradation of oxidatively truncated PCs (oxPCs) generated by oxidative stress as well as medium-chain phospholipids[27]. It has been suggested that heat alters biomembranes[7]. Heat can also affect mitochondria, which are cellular organelles, and may contribute to the production of ROS[5]. *ABHD3* may contribute to biomembrane homeostasis by removing oxPCs generated by ROS during HS. In contrast, we found that *ZFAND2A* and *USPL1* had high ChIP-seq scores for the three transcription factors (Fig. 5A-C). *ZFAND2A* is a gene whose expression is induced by heat, and HSF1 has been suggested to be involved in this process[28]. Notably, there have been few reports of this gene, despite reports of its association with HS. These examples indicate that HS research focuses on the expression and functional analysis of molecular chaperones. However, there is no current evidence suggesting that *USPL1* is associated with HS. It has been suggested that *USPL1* encodes a protein that does not target ubiquitin but functions as a small ubiquitin-related modifier (SUMO) isopeptidase[29]. However, according to gene2pubmed data, there have been very few reports from human (20 articles published as of April 2023), and its functions still need to be fully understood. The list of genes obtained in this study featured many genes, including *USPL1*, whose role in HS is yet to be fully elucidated. The analysis of such genes may contribute to a better understanding of the overall response to HS, thus warranting further research. Further functional analysis should include the use of genome editing tools, such as CRISPR-Cas9, which enable precise manipulation of target genes and are especially useful for the functional analysis of genes whose functions are unknown for the HS extracted in this study. Therefore, the genes identified in this study may serve as candidates for novel genome-editing target genes.

In this study, a meta-analysis of human and mouse gene expression data facilitated the extraction of genes not yet known to be associated with HS, in addition to the responses of organisms to heat derived from previous findings. Such findings from data-driven studies can potentially provide novel insights. However, it is unclear whether the genes identified in this study are as widely conserved in species as molecular chaperone genes; therefore, a meta-analysis of gene expression data in a broader range of species should be conducted to identify novel common mechanisms of HS responses. This knowledge could contribute to addressing the increased exposure to and frequency of high temperatures due to climate change. With growing concerns about increased health hazards and impacts on industries, such as crop production[1], this research lays the groundwork for developing countermeasures to these problems and may lead to new knowledge and technological advances in future research.

## Methods

### Curation of gene expression data from public database

The public database, Gene Expression Omnibus (GEO)[30], was used to obtain gene expression data associated with human and mouse HS. GEO is operated by the National Center for Biotechnology Information (NCBI) and archives gene expression data obtained by RNA sequencing (RNA-seq) using Next-Generation Sequencing (NGS) and expression microarrays. To narrow down the gene expression data, we used keywords related to heat exposure such as “heat stress,” “heat shock,” “hyperthermia,” “thermal stress,” “heat-shock treatment,” and “heat stroke.” Additionally, “Expression profiling by high throughput sequencing” was added to the search formula as a Study type condition. Data collection was performed manually, allowing for pairs of HS and non-treatment data and the description of data such as temperature conditions and sample type.

### Quantification of gene expression data using the analysis pipeline

Gene expression data retrieval, processing, and quantification were performed using ikra (ver. 2.0.1)[31], an automated human and mouse RNA-seq data analysis pipeline. ikra is a tool that can automatically perform all the following processes: retrieval of FASTQ format files using the fastq-dump program in the NCBI SRA tool kit (ver. 3.0.0)[32], read quality control, and adapter trimming using Trim_galore (ver. 0.6.6)[33] and transcript quantification using Salmon (ver. 1.9.0)[34]. This study used a different version of the tool compared to that used in previous studies[16,17]: the SRAtoolkit, Salmon. Different versions of reference sequence sets used as indices in salmon were also used, including GENCODE Release 37 (GRCh38.p13) in human and GENCODE Release 26 (GRCm39) in mouse.

### Calculation of HN-ratio and HN-score

Expression ratios (henceforth referred to as the HN-ratio) were calculated for all genes from gene expression data paired with HS and non-treatment. The HN-ratio for each gene was calculated using the following equation (1):

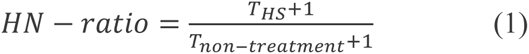

*T_HS_* and *T_non-treatment_* refer to scaled Transcripts Per Million (scaled TPM)[35] under HS and non-treatment conditions, respectively, and indicate the quantified expression levels. When calculating the HN-ratio, 1 was added to the expression value to avoid calculation with a zero value[17,36].

### Classification of genes based on HN-ratio

Using the calculated HN-ratio, all genes were classified into three groups: upregulated, downregulated, and unchanged. If the HN-ratio was higher than a threshold, the gene was considered “upregulated.” Conversely, if the HN-ratio was less than the reciprocal of a threshold, the gene was considered “downregulated.” Genes not classified in either group were considered “unchanged.” For genes that were upregulated, we tested 2-fold, 5-fold, and 10-fold thresholds and selected the 5-fold threshold; for genes that were downregulated, we tested 1/2-fold, 1/5-fold, and 1/10-fold thresholds and selected the 5-fold threshold for classification.

### Calculation of HN-score

To evaluate the genes whose expression was altered by HS, an index called the Heat-stress and Non-treatment score (HN-score) was calculated for each gene in human and mouse. The HN-score was calculated by subtracting the number of instances of genes classified as downregulated from those of genes classified as upregulated. The HN-ratio and HN-score were calculated using a script from a previous study (https://github.com/no85j/hypoxia_code) [16]. Python scripts were created to visualize the HN-score values for each gene using a scatter plot (https://github.com/yonezawa-sora/HS_code).

### Analysis of selected gene sets in human and mouse

From each human and mouse dataset, the top and bottom genes were selected based on the HN-score, and gene set enrichment analysis was performed. Gene set enrichment analysis was performed using the web tool Metascape[37] (accessed November 2022). The Batch Query function of mouse genome informatics[19] (accessed November 2022) was used to extract genes that are commonly conserved between human and mouse. Scatter plots of the HN-scores of human genes corresponding to mouse “Current gene symbol” and the HN-scores of mouse genes were created using a Python script. The scripts used are available on GitHub (https://github.com/yonezawa-sora/HS_code). Gene set enrichment analysis, protein–protein interaction networks, and transcription factor target gene set enrichment analyses were performed using Metascape[37]. Protein– protein interaction networks were processed using Cytoscape[38]. Genome mapping was performed using ShinyGO[20] (ver. 0.77) (accessed February 2023). A Python script was used to create a stacked bar graph to visualize the contribution of each sample to the HN-score. The script is available on GitHub (https://github.com/yonezawa-sora/HS_code).

### Comprehensive analysis of common upregulated genes in human and mouse

The ChIP-Atlas Database[22] (accessed February 2023) was accessed, and the “Target Genes” function was used to obtain an average model-based analysis of ChIP-seq (MACS2) scores for all genes. HSF1, HSF2, and PPARGC1A (PGC1-α) were used as “antigens.” and MACS2 scores were obtained at distances of ± 5 kb from the transcription start site. The MACS2 scores for genes not listed were 0. For each gene, the calculated HN-score, MACS2 score, and number of literature reports for the human gene in gene2pubmed data (accessed February 2023) were visualized by creating a scatter plot using a Python script. The script used for the visualization is available on GitHub (https://github.com/yonezawa-sora/HS_code).

## List of abbreviations

HS: Heat stress
HSP: Heat shock protein
HSF: Heat shock factor
HN-ratio: Heat stress-non-treatment ratio
HN-score: Heat stress-non-treatment score

## Declarations

### Ethics approval and consent to participate

Not applicable

### Consent for publication

Not applicable

### Availability of data and materials

The data presented in this study are publicly available in figshare[18].

All the scripts used in this study are publicly available on GitHub (https://github.com/yonezawa-sora/HS_code).

### Competing interests

The authors declare that they have no competing interests.

## Funding

This work was supported by the Center of Innovation for Bio-Digital Transformation (BioDX), an open innovation platform for industry–academia cocreation (COI-NEXT) and the Japan Science and Technology Agency (JST, COI-NEXT, JPMJPF2010).

### Authors’ contributions

Conceptualization, S. Y. and H. B.; methodology, S. Y. and H. B.; software, S. Y. and H. B.; validation, S. Y. and H. B.; formal analysis, S. Y. and H. B.; investigation, S. Y.; resources, S. Y. and H. B.; data curation, S. Y.; writing—original draft preparation, S. Y.; writing—review and editing, S. Y. and H. B.; visualization, S. Y.; supervision, H. B.; project administration, H. B.; funding acquisition, H. B. All authors read and agreed to the published version of the manuscript.

## Acknowledgements

Computations were performed on the computers at Hiroshima University Genome Editing Innovation Center.

## Notes

https://doi.org/10.6084/m9.figshare.c.6564487

### Competing Interest Statement

The authors have declared no competing interest.

