## Supplemental Figures for "Meta-analysis of heat-stressed transcriptomes using the public gene expression database from human and mouse samples"

Fig. S1

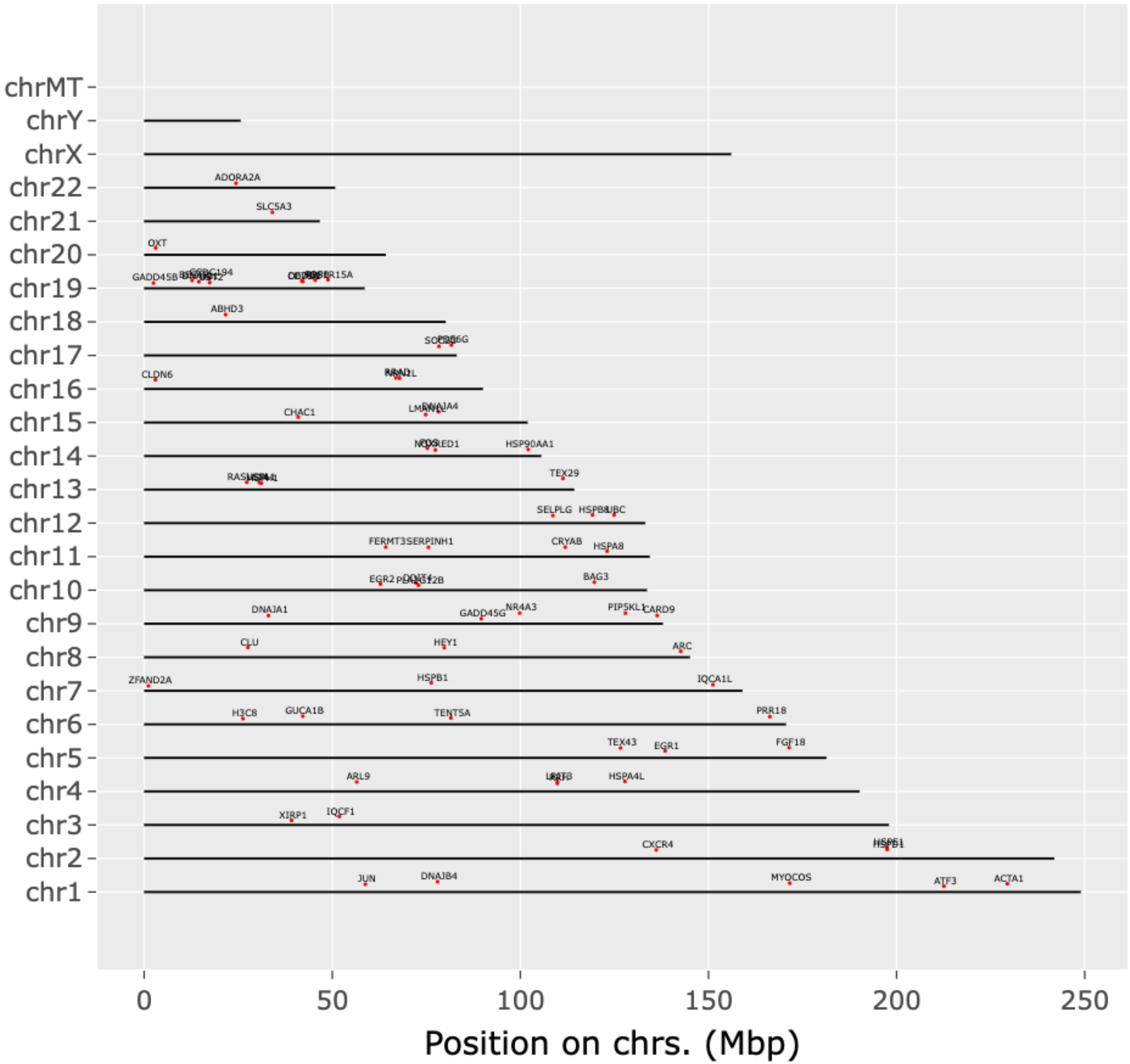

chrMT
chrY
chrX
chr22
chr21
chr20
chr19
chr18
chr17
chr16
chr15
chr14
chr13
chr12
chr11
chr10
chr9
chr8
chr7
chr6
chr5
chr4
chr3
chr2
chr1
ADORA2A
SLC5A3
OXT
GADD45B
ABHD3
CLDN6
CHAC1
DNAJA4
HSP90AA1
TEX29
SELPLG
UBC
FERMT3
SERPINH1
CRYAB
HSPA8
EGR2
BAG3
DNAJA1
GADD45G
NR4A3
CARD9
CLU
HEY1
ARC
ZFAND2A
HSPB1
IQCA1L
H3C8
GUCA1B
TENT5A
PRR18
TEX43
EGR1
FGF18
ARL9
HSPA4L
XIRP1
IQCF1
CXCR4
JUN
DNAJB4
MYOCOS
ATF3
ACTA1
0
50
100
150
200
250
Position on chrs. (Mbp)

Fig. S1

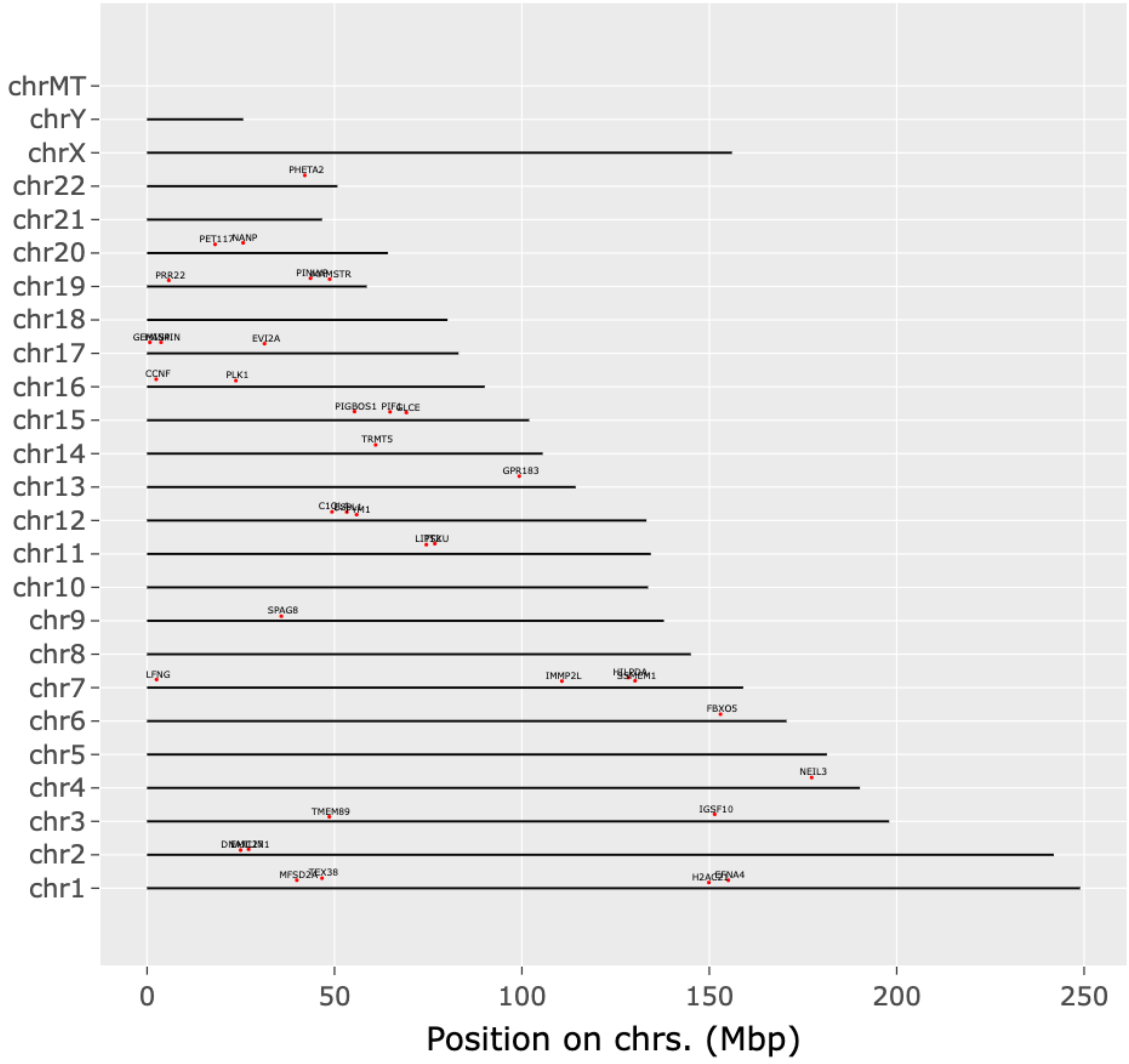

chrMT
chrY
chrX
chr22
chr21
chr20
chr19
chr18
chr17
chr16
chr15
chr14
chr13
chr12
chr11
chr10
chr9
chr8
chr7
chr6
chr5
chr4
chr3
chr2
chr1
PHETA2
NANP
PRR22
EVI2A
CCNF
PLK1
TRMT5
GPR183
SPAG8
LFNG
IMMP2L
FBXO5
NEIL3
TMEM89
IGSF10
MFSD2A
TEX38
0
50
100
150
200
250
Position on chrs. (Mbp)

Fig. S2

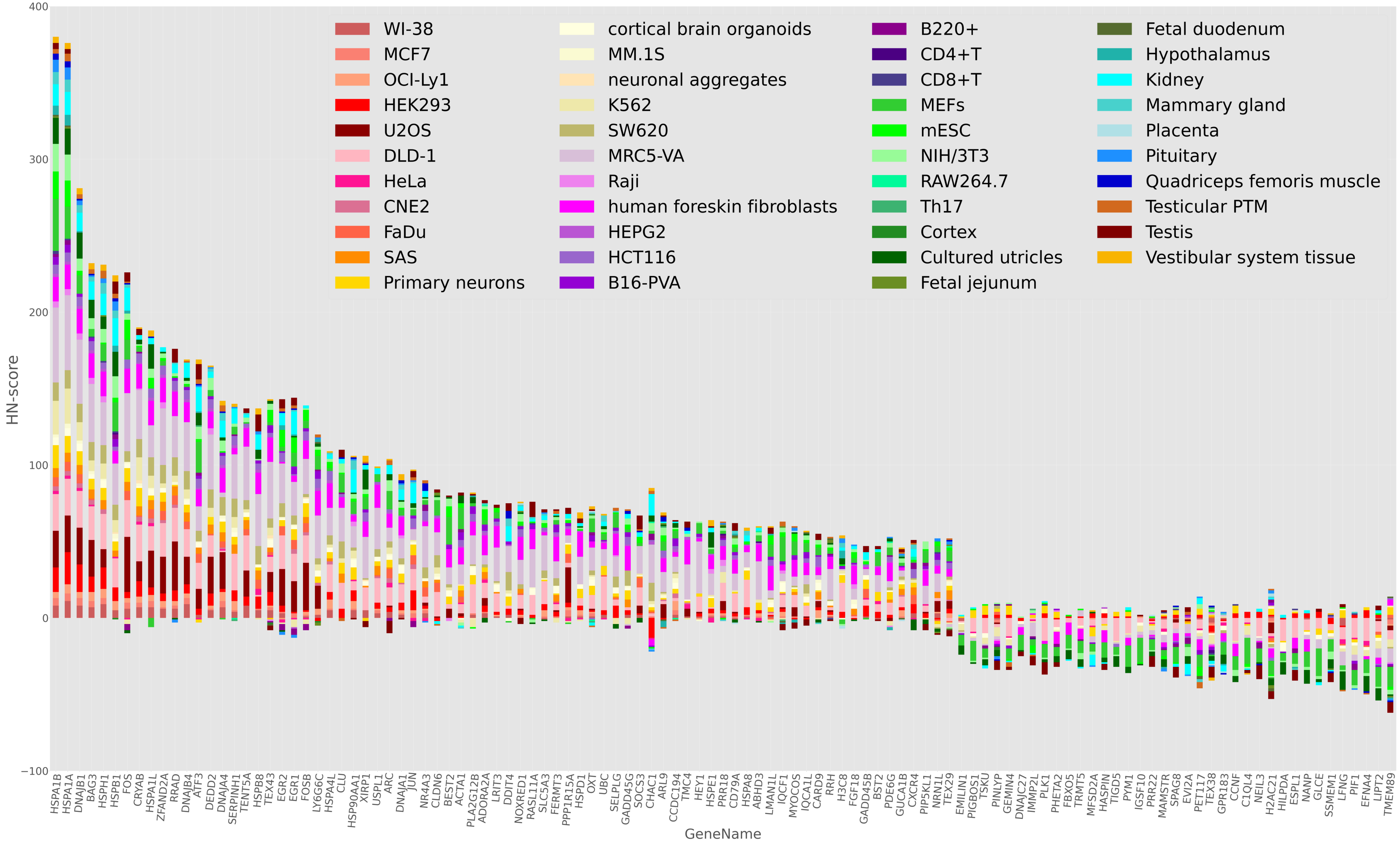
WI-38
MCF7
OCI-Ly1
HEK293
U2OS
DLD-1
HeLa
CNE2
FaDu
SAS
Primary neurons
cortical brain organoids
MM.1S
neuronal aggregates
K562
SW620
MRC5-VA
Raji
human foreskin fibroblasts
HEPG2
HCT116
B16-PVA
B220+
CD4+T
CD8+T
MEFs
mESC
NIH/3T3
RAW264.7
Th17
Cortex
Cultured utricles
Fetal jejunum
Fetal duodenum
Hypothalamus
Kidney
Mammary gland
Placenta
Pituitary
Quadriceps femoris muscle
Testicular PTM
Testis
Vestibular system tissue
HN-score
400
300
200
100
0
−100
GeneName
HSPA1B
HSPA1A
DNAJB1
BAG3
HSPH1
HSPB1
FOS
CRYAB
HSPA1L
ZFAND2A
RRAD
DNAJB4
ATF3
DEDD2
DNAJA4
SERPINH1
TENT5A
HSPB8
TEX43
EGR2
EGR1
FOSB
LY6G6C
HSPA4L
CLU
HSP90AA1
XIRP1
USPL1
ARC
DNAJA1
JUN
NR4A3
CLDN6
BEST2
ACTA1
PLA2G12B
ADORA2A
LRIT3
DDIT4
NOXRED1
RASL11A
SLC5A3
FERMT3
PPP1R15A
HSPD1
OXT
UBC
SELPLG
GADD45G
SOCS3
CHAC1
ARL9
CCDC194
TMC4
HEY1
HSPE1
PRR18
CD79A
HSPA8
ABHD3
LMAN1L
IQCF1
MYOCOS
IQCA1L
CARD9
RRH
H3C8
FGF18
GADD45B
BST2
PDE6G
GUCA1B
CXCR4
PIP5KL1
NRN1L
TEX29
EMILIN1
PIGBOS1
TSKU
PINLYP
GEMIN4
DNAJC27
IMMP2L
PLK1
PHETA2
FBXO5
TRMT5
MFSD2A
HASPIN
TIGD5
PYM1
IGSF10
PRR22
MAMSTR
SPAG8
EVI2A
PET117
TEX38
GPR183
CCNF
C1QL4
NEIL3
H2AC21
HILPDA
ESPL1
NANP
GLCE
SSMEM1
LFNG
PIF1
EFNA4
LIPT2
TMEM89

Fig. S3

(A)

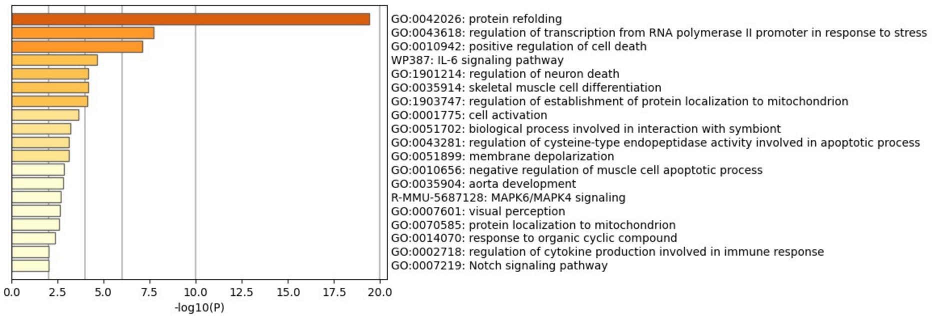


(B)

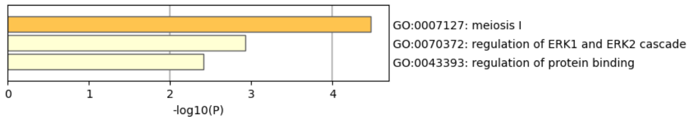


(C)

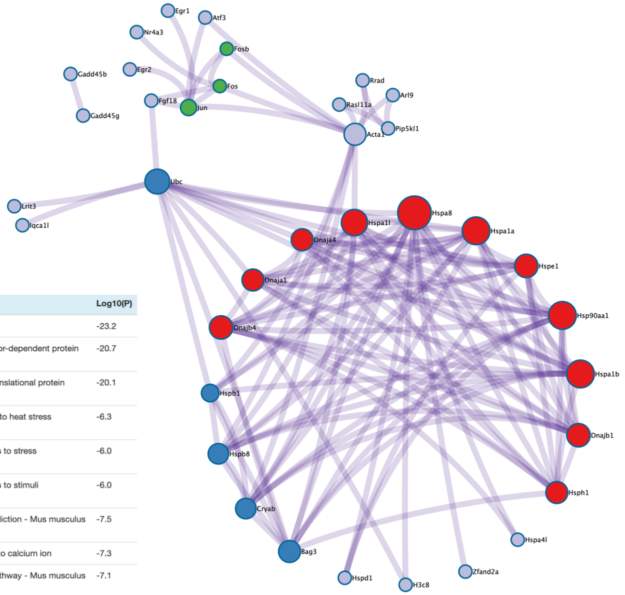


| Color | MCODE | GO | Description | Log10(P) |
|---|---|---|---|---|
| red | MCODE_1 | GO:0006457 | protein folding | -23.2 |
| red | MCODE_1 | GO:0051085 | chaperone cofactor-dependent protein refolding | -20.7 |
| red | MCODE_1 | GO:0051084 | 'de novo' post-translational protein folding | -20.1 |
| blue | MCODE_2 | R-MMU-3371556 | Cellular response to heat stress | -6.3 |
| blue | MCODE_2 | R-MMU-2262752 | Cellular responses to stress | -6.0 |
| blue | MCODE_2 | R-MMU-8953897 | Cellular responses to stimuli | -6.0 |
| green | MCODE_3 | mmu05031 | Amphetamine addiction - Mus musculus (house mouse) | -7.5 |
| green | MCODE_3 | GO:0071277 | cellular response to calcium ion | -7.3 |
| green | MCODE_3 | mmu04657 | IL-17 signaling pathway - Mus musculus (house mouse) | -7.1 |

(D)

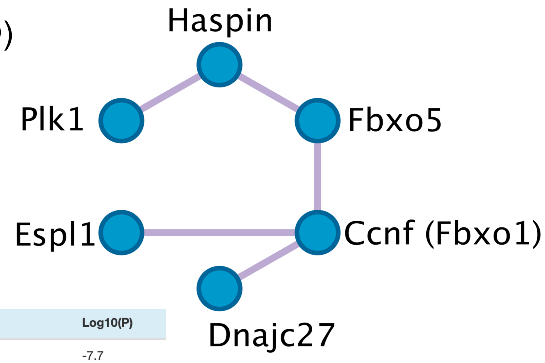


| GO | Description | Log10(P) |
|---|---|---|
| GO:0007127 | meiosis I | -7.7 |
| GO:0061982 | meiosis I cell cycle process | -7.7 |
| GO:0140013 | meiotic nuclear division | -7.1 |

Fig. S4

(A) Common upregulated genes transcription factor target

(B) Upregulated genes Upset plots

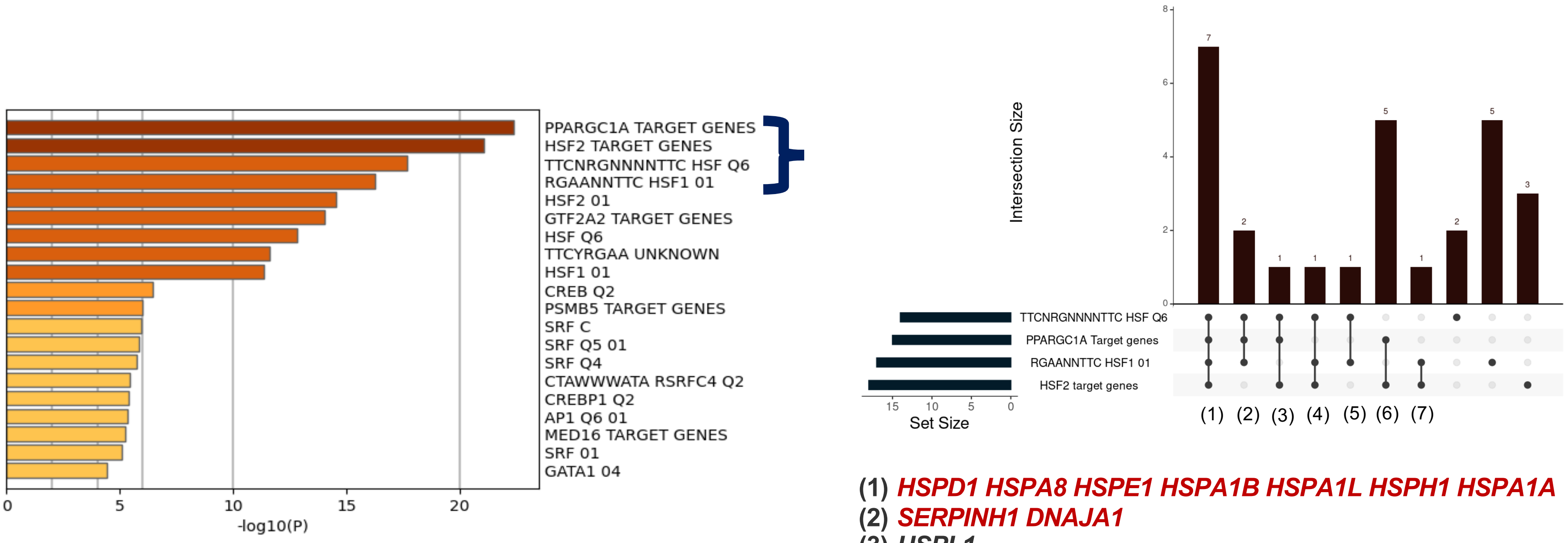


(1) *HSPD1 HSPA8 HSPE1 HSPA1B HSPA1L HSPH1 HSPA1A*
(2) *SERPINH1 DNAJA1*
(3) *USPL1*
(4) *CRYAB*
(5) *NR4A3*
(6) *BAG3 HSP90AA1 ZFAND2A SLC5A3 BST2*
(7) *ABHD3 HSPB1*
